# Salt-bridge Networks within Globular and Disordered Proteins – Characterizing Trends for Designable Interactions

**DOI:** 10.1101/113621

**Authors:** Sankar Basu, Debasish Mukharjee

**Affiliations:** Department of Biochemistry, University of Calcutta 35, Ballygunge Circular Rd, Ballygunge, Kolkata, West Bengal 700019, India; Computational Science Division, Saha Institute of Nuclear Physics 1/AF, BidhanNagar, Kolkata, West Bengal 700064, India

**Keywords:** salt-bridges, ioinic bonds, electrostatic complementarity, motif identifier, bifurcation angle, bifurcated salt-bridges, Molecular clips, Intrinsically Disordered Proteins

## Abstract

There has been fare amount of debate regarding the contribution of salt-bridges in the stabilization of protein folds. However, their participation in crucial protein functions are well established. The current study analyzes their modes of association, in terms of networks, both within globular proteins and also at protein-protein interfaces. Apart from the most common and trivial case of isolated salt-bridges, bifurcated salt-bridges appear to be a special salt-bridge motif both in terms of its topology and geometry and found ubiquitously in proteins and inter-protein complexes. Interesting and attractive examples presenting different interaction-modes have been highlighted. Bifurcated salt-bridges appear to function as molecular clips instrumental in stitching large surface contours of interacting protein-protein interfaces. The work also emphasizes the key role of salt-bridge mediated interactions in the partial folding of proteins containing large amount of disordered regions. Salt-bridge mediated interactions seem pivotal in promoting ‘disorder-to-order’ transitions for small disordered protein fragments and their stabilization upon binding. The results should guide to elucidate the modus operandi of these partially disordered proteins and also should be helpful to conceptualize how these proteins manage to keep necessary amount of disorder even in their functionally active bound forms, encouraging future studies. It should also be potentially beneficial towards the proposed notion of geometrically specific designable interactions involving salt-bridges.

## Introduction

Salt-bridges were envisaged to be instrumental primarily in sustaining the global electrostatic balance within correctly folded globular proteins and also at protein-protein interfaces [1], [2]. However, that theory had to be revised and rephrased as only partially true at least in the context of protein-protein interfaces [3] after the advent of continuum electrostatic models which could numerically solve the Poisson-Boltzmann equation for the protein-solvent system [4], [5]. It was categorically shown that electrostatic complementarity (EC) at the protein-protein interfaces could still retain significantly high values, even when the salt-bridges were computationally neutralized (or switched off). The reason for this is that the electrostatic complementarity is a non-local effect governed by a long-range force, wherein partial charges coming from all atoms from the two interacting molecules are collectively contributing to the potentials, and, hence, switching off the salt-bridges did not really alter the overall correlation. In fact, this was the very first exclusive study on electrostatic complementarity at protein-protein interfaces [3] wherein the earlier idea of charge complementarity was virtually ruled out and replaced by the concept of complementarity in terms of electrostatic potential. That is to say that the complementarity is actually attained in the anti-correlation of surface electrostatic potentials realized at the protein-protein interfaces. A later study extended this concept to the folding of single domain globular proteins and for the first time, characterized the trends in electrostatic complementarity in a residue-wise manner within the protein interior [6]. In consistency to that of the protein-protein interfaces, here also, elevated values in electrostatic complementarity (E_m_) was attained for all completely or partially buried amino acid residues, irrespective of their chemical identity, both, in presence and absence of the contribution of the salt-bridges. All said and done, the different functions attributed to these ionic bonds ranging from anchoring protein subunits [7] to metal coordination and co-operativity in networks of salt-bridges [8] as well as their impact on stability [9], [10] can not be undermined.

There does not appear to be a universal rule regarding the role of salt-bridges in stabilizing protein structures though. Due to desolvation effects, they are generally considered to be destabilizing [11], though instances have been observed where networks of ionic bonds contribute favorably to the thermal stabilization of proteins [10], [12], [13]. For instances, arginine-glutamate isolated salt-bridges have been found to be favorably contributing to the stability of helices [10], whereas bifurcated salt-bridges have been shown to be instrumental in originating photoacoustic signals in cytochromes [14].

Although some empirical rules have been proposed regarding the geometry of hydrogen bonds to be found within salt-bridges [7], [13], and also the distinction between simple and complex salt-bridges [7], a comprehensive characterization of different possible unique topological assemblies of salt-bridge networks consistent with the growing number of high resolution protein structures was felt lacking.

In this background, the current study presents an exclusive characterization of salt-bridge networks within proteins in terms of their topology and geometry and composition. The study further recognizes bifurcated salt-bridges as a key motif found in prevailing majority within proteins, both at the interior and at interfaces. Along that line, the study constitutes a statistical analyses on the functional attributes of bifurcated salt-bridges, put forward an exhaustive list of functionally relevant interactions and highlights some unique and interesting patterns. It also attempts to shed some light on the potential contribution of ionic bonds and charge residue contacts in the partial folding / stabilization of intrinsically disordered proteins (IDPs) both in their free and bound forms. The study should be beneficial in the context of designing salt-bridges in recombinant proteins and also in the computational modeling of interaction involving disordered protein regions.

## Materials and Methods

### Databases

#### Globular Proteins

Following our previous reports in the related area of protein electrostatics [2], we went on accumulating salt-bridges from a previously reported [6], [15] database (https://github.com/nemo8130/DB2) of high resolution crystal structures of native globular proteins. The structures in the culled database (resolution ≤ 2Å; R-factor ≤ 20%; pair-wise sequence similarity ≤ 30%) were devoid of any bulky prosthetic groups (e.g., protoporphyrins) and any missing atoms. This database was used to characterize the topological and geometric preferences of networks involving salt-bridges.

#### Protein-protein complexes

To study the role of salt-bridges in triggering and stabilizing protein-protein interactions, another database containing high resolution (≤ 2Å) crystal structures of 1879 native protein-protein complexes was assembled. None of these structures contained any missing backbone atoms. This database was previously used in structure validation of protein complexes [16] and also as a reality check in protein-protein docking scoring [17]. In case of structures containing more than two chains, the two largest interacting chains were considered for all subsequent calculations.

#### Proteins with Intrinsically Disordered Regions

To study the role salt-bridges and atomic contacts involving charged residues in the stabilization of intrinsically disordered proteins, yet another non-redundant database of 109 polypeptide chains was assembled containing missing disordered loops. This database was a subset of an originally assembled set and used in a recent study on conformational entropy of intrinsically disordered proteins calculated from amino acid triads [18]. The original database [18] contained all types of possible experimental structures solved by X-ray Crystallography, NMR and Electron Microscopy. Missing disordered residues were identified by comparing the SEQRES and ATOM records in the PDB files and then cross-checking the difference between the two sets with the missing residues declared in the REMARK 465 list. The database was initially compiled using the PISCES server [19] with the selection criteria of sequence similarity ≤ 25% and length ≥ 40 residues resulting in a total of 138 chains with > 50% structural disorder [18]. Over and above the selection criteria implemented previously, we further discarded the EM-structures and structures containing more than 700 residues reducing it to 109 chains.

Apart from the systematic calculation performed on the above database, two case studies were also presented on two popular individual systems involving disorder. The first of the two was the most popular Tau protein (PDB IDs: 2N4R, 4TQE, 5DMG and 5BTV), one of the potential causal factors in the Alzheimer’s disease due to its abnormal phosphorylation resulting in paired helical filament and neurofibrillary tangles. The second candidate was the plant protein Crambin solved at an ultrahigh resolution having the special feature of valance electron density (PDB ID: 1EJG) presenting an unique case with all isoforms being individually disordered (http://deposit.rcsb.org/format-faq-v1.html).

### Modeling IDPRs in existing experimental structures

Disordered residues were identified and distinguished by the selection procedure elaborated in the database section. The missing disordered residues (as listed in the REMARK 465 of the corresponding PDB file) were then built by MODELLER (version 9.17) [20] implementing its general ‘automodel’ or advanced ‘loopmodel’ modules (the latter with the ‘fast refinement’ option) as appropriate to each individual protein system. The experimental structure was used as a template for modeling and the correct position of each missing residue was determined by a pairwise sequence alignment performed using the high-accuracy alignment program MUSCLE (version 3.8) [21] and fed to MODELLER. The root mean square (rms) deviation calculated from all pair of corresponding atoms (i.e., the atoms solved experimentally) upon aligning the two structures were found to be ≤ 1.5 Å for all models. For each case the appropriate MODELLER module (automodel / loopmodel) was determined by a comparative study of the two initial models produced by the two methods and the one resulting in the lowest rms deviation was chosen. Once the correct module was determined for each protein, the program was rerun generating 10 models and the one with the lowest rms deviation upon alignment to the corresponding ‘experimental’ template was chosen (**Fig.1**). Residues in the modeled structures were then renumbered (from 1) according to the SEQRES records to naturally accommodate the missing disordered residues avoiding negative integers and reported that exact way. Stride [22] was used to calculate the relative content of secondary structural elements in these modeled structures.

**Figure 1.**
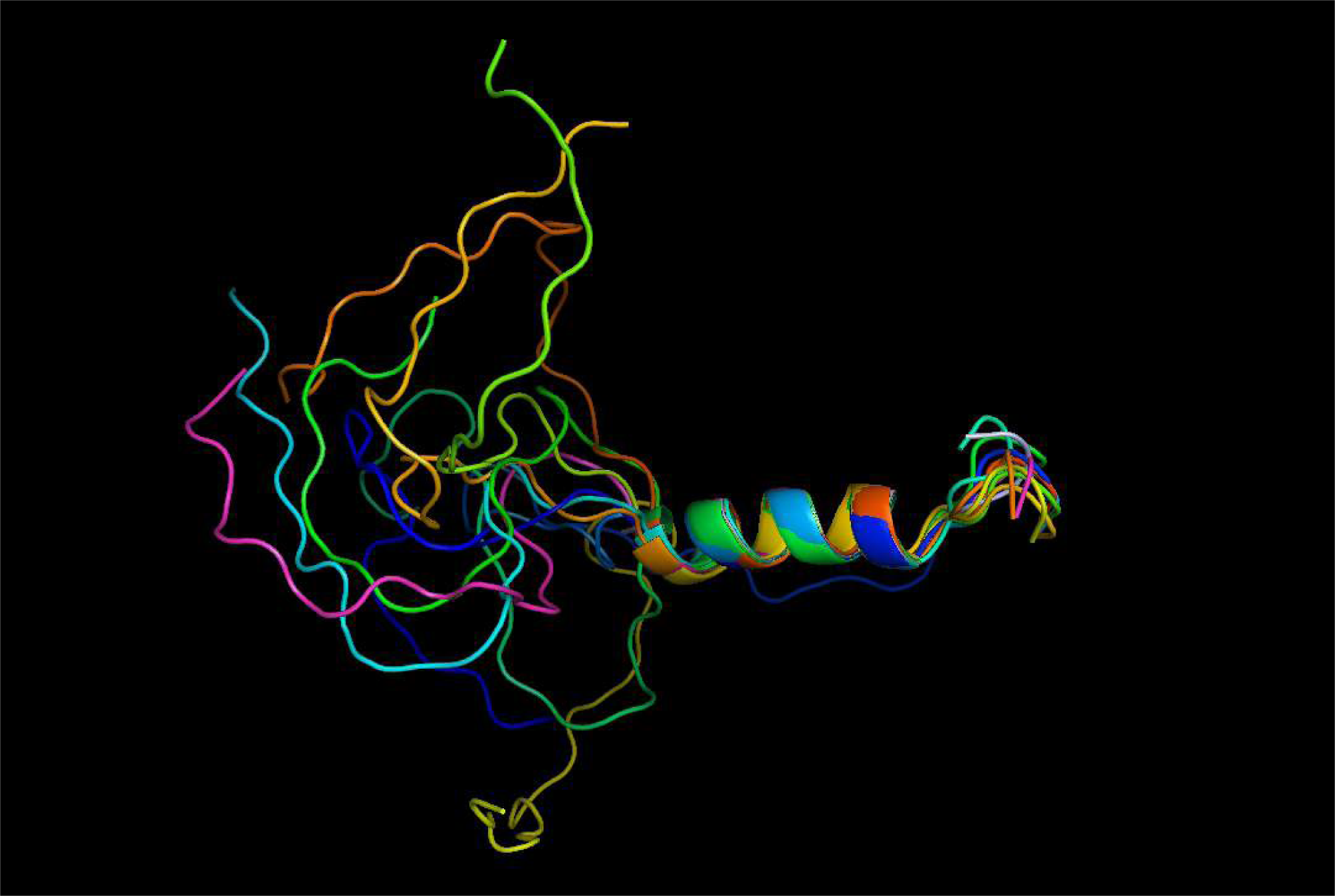
Modeling IDPRs within partially disordered proteins. The example shows top 10 models created by the ‘loopmodel’ module of MODELLER for the partially disordered protein 2MBH modeling its missing IDPRs. All models were closely spaced in terms of agreeing with the native experimental template and gave rise to rms deviations (calculated from all experimentally solved atoms) ranging from 0.461 A to 1.018 A upon superposition. The native and the best model (rmsd: 0.461 Å) have been colored by light-blue and light-magenta respectively.

### Burial of Solvent Exposure

Solvent accessible surface area (ASA) was calculated for each atom in a protein molecule by NACCESS [23] by rolling a probe sphere of 1.4 Å over the entire protein surface (the Lee and Richards algorithm) [24]. The ASAs were then summed up for all atoms pertaining to the same residue. This was followed by calculating the burial of their solvent exposure defined as the ASA of a residue, X located in the protein divided by the ASA of the same amino acid in a Gly-X-Gly peptide fragment in its fully extended conformation following a previously established method [25]. The completely exposed residues having a residue burial > 0.3 [25] were then identified and the atoms demarcated as the peripheral atoms of the protein.

### Accessibility Score

Depending on the location and extent of disorder embedded in a protein sequence, the modeled protein can in principle be delirious and unreal. One way to estimate their fury is to check the expected distribution of amino acid residues with respect to their burial of solvent exposure and compare the values with globular proteins known to be stable in solutions. This was estimated by the accessibility score, *rGb* which have previously been standardized both within globular proteins [15] and at protein-protein interfaces [16] and have also been successfully used as a coordinate driven feature in protein-protein docking scoring [17]. Briefly, the definition of the *rGB* score is as follows.

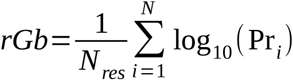

where N_res_ is the total number of residues in the polypeptide chain and Pr_i_ is the propensity of a particular amino acid (Val, Leu, Asp etc) to acquire a particular degree of solvent exposure. Other details of the score is defined elsewhere [15]. The *rGb* score raised similar range of values for both the correctly folded globular proteins (0.055 ±0.022) as well as the protein-protein interfaces (0.058 ±0.022). Based on these observations, its upper threshold characteristic of either a well-folded protein or a well-packed interface was determined to be 0.011. In principle, the score is also applicable to peptides and a lower value than the threshold (which can also be negative) is indicative of instability of the protein / peptide in solution and such molecules will be highly reactive.

### Defining Salt-bridges and their Networks

The essential idea behind the analyses on the database of globular proteins was to collect salt-bridges not merely as a collection of individual ionic bonds, rather as networks, and perform a exhaustive statistical study on the topology and geometrical preferences of different salt-bridge networks. In these networks, charged residues were represented as nodes and connected by an edge when there exist an ionic bond (or salt-bridge) between them. The ionic bonds were detected when a positively charged nitrogen atom of lysine (NZ), arginine (NH1, NH2) or positively charged histidine (HIP: ND1 NE2, both protonated) were found to be within 4.0 Å of a negatively charged oxygen atom of glutamate (OE1, OE2) or aspartate (OD1, OD2).

In order to study the topological and geometric patterns of these networks, constituted by ionic bonds, a previously proposed numeric scheme, namely, the motif identifier was adapted [26] which could be considered as a reduced representation of a specific unique network topology. In this numerical representation, an unique network-topology is represented by *n* concatenated strings of numbers separated by delimiters, where, *n* is the number of nodes in the network. Each string begins with the degree of a node (from the highest degree node following a descending order in degrees), followed by the degrees of its linked nodes again sorted in descending order (**Fig.2**). The whole idea and fromulation of the numeric topological scheme was based on the assumption that a node in such networks is always limited in its number of direct neighbors, (due to steric and electrostatic constraints) to single-digit numbers (i.e., a maximum of 9) – which was found generally true in a wide plethora of protein contact networks [26].

**Figure 2.**
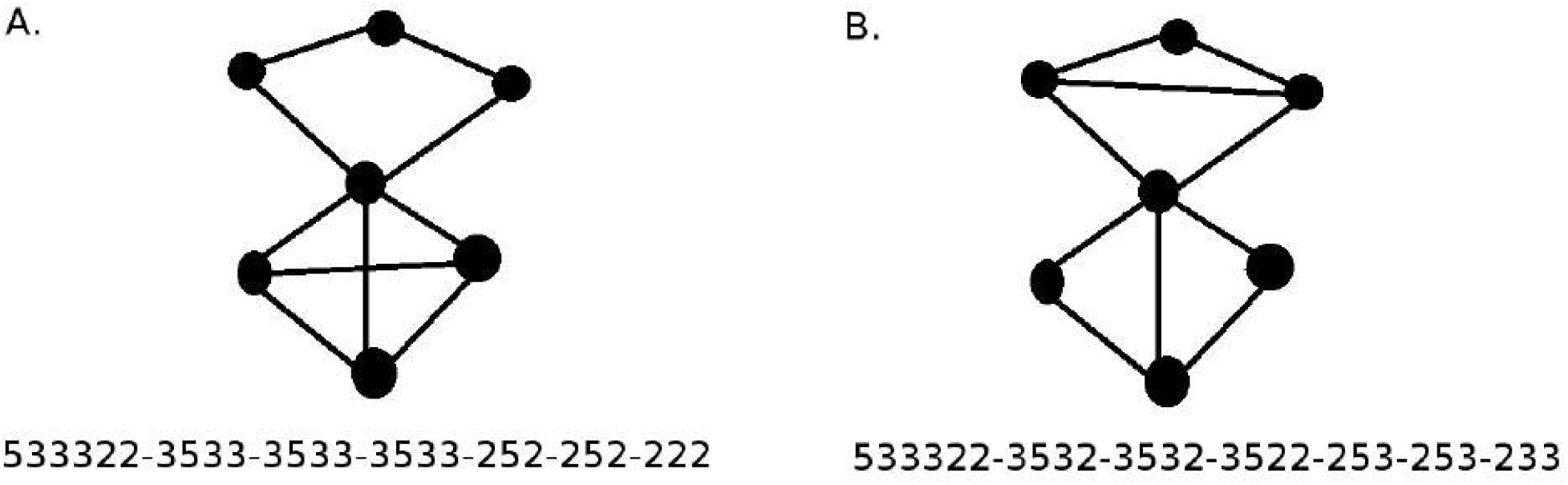
The motif identifier: accounting for topological variations in graphs. The motif identifier (presented as the concatenated sorted numeric string) is a collection of numeric sub-strings each of which is representative of a topological motif pertaining to a node (nodal motif). The first number in each numeric string stands for the degree of the corresponding node and the other numbers represent the degrees of its direct neighbors sorted in a descending order. All nodal motifs hence obtained are again collected as elements of an array and further sorted in descending order. Finally the sorted number strings (nodal motifs) are concatenated using a delimiter (‘-’). Two distinct examples have been shown to demonstrate the numeric scheme. Both graphs are generated starting from a common core backbone topology and the rewiring a few links.

Thus, when put to perspective, the motif identifier essentially discriminates between two graphs based on the combined distribution of degrees of their constituent nodes coupled with the degrees of their neighboring nodes; and, will potentially signal for any variability in these network parameters between two given graphs (**Fig.2**). In principle, the only instance where it fails to discriminate between two non-identical networks is the case of k-regular graphs [27] with k>2. Since for regular graphs, all nodes have identical degrees, no variability could be accounted for in terms of their degree and/or the degrees of their neighboring nodes, even if the two k-regular graphs of the same size are topologically non-identical. However, such graphs are irrelevant in the context of salt-bridges due to steric and electrostatic constraints.

### Amino acid Propensities to form salt-bridges

Propensity (*Pr(x,s)*) of a charged residue, *x* to go into a salt-bridge was computed by the following expression:

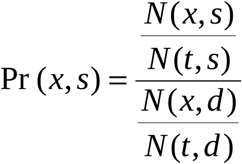

where *N(x,s)* is the count of the residue *x* found in salt-bridges, *N(t,s)* is the total number of residues involved in salt-bridges, *N(x,d)* and *N(t,d)* are the counts of residue *x* and the total number of residues in the database.

### Electrostatic Complementarity of charged amino acids

The electrostatic complementarity (*E_m_*) for charged amino acid residues within the protein interior was calculated according to a previous report [6] and a detailed residue wise statistics were performed on them. To start with, hydrogen atoms were geometrically fixed to each individual protein chain in the database. Van der Waals surfaces were sampled at 10 dots / Å^2^. The details of the surface generation have been discussed elsewhere [25]. The exposure of individual atoms to solvent was estimated by rolling a probe sphere of radius 1.4 Å over the protein atoms [24] and the burial of individual residues was estimated by the ratio of solvent accessible surface areas of the charged amino acid X embedded in the polypeptide chain to that of an identical residue located in a Gly–X–Gly peptide fragment with a fully extended conformation. Partial charges and atomic radii for all protein atoms were assigned in consistency with a previously used force field [28] and Asp, Glu, Lys, Arg, doubly-protonated histidine (Hip) along with both the carboxy, amino terminal groups were considered to be ionized / charged. Crystallographic water molecules and surface bound ligands were excluded from the calculations and thus modeled as bulk solvent. Ionic radii were assigned to the bound metal ions according to their charges [29].

Delphi (version 6) [30] was used to compute the electrostatic potential of the molecular surface along the polypeptide chain according to a previous report [6]. For each run, a set of dot surface points on which the electrostatic potentials are to be computed were fed to Delphi along with a set of charged atoms contributing to the potential. First, the dot surface points of the individual amino acids (targets) were identified. The electrostatic potential for each residue surface was then calculated twice, 1) due to the atoms of the particular target residue and 2) from the rest of the protein excluding the selected target amino acid. In either case, the atoms not contributing to the potential were considered as dummy atoms with only their radii being assigned with zero charge. Thus each dot surface point of the target residue was then tagged with two values of electrostatic potential. The electrostatic potential complementarity **(*E_m_*)** of an amino acid residue (at the protein interior) was then defined as the negative of the Pearson’s correlation coefficient between these two sets of potential values [6].

Again, the potential values corresponding to N dot surface points could be divided into two distinct sets, based on whether the dot point is sourced from main-or side-chain atoms of the target residue and ***E**_m_* could be obtained separately for each set. In the current study, however, we were only interested in the ***E**_m_* obtained from the side-chain dot surface points (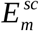) since its the side-chain atoms which bear the charges, in the context of ionic bonds.

All images representing protein structures were created in Pymol (The PyMOL Molecular Graphics System, Version 1.8 Schrödinger, LLC.) for visual investigation and display.

## Results and Discussion

### Characterizing Salt-bridge Networks within proteins

A total of 3076 networks were extracted from the globular protein database using the numerical scheme described in the previous section (see **Materials and Methods**). The distribution of such networks was found to be dominated by isolated ionic bonds (**11-11:** 2445, **Fig.3**) followed by bifurcated salt-bridges consisting of three nodes (**211-12-12:** 475). For networks with number of nodes greater than three, the overwhelming majority fell into the class of open linear chains, or, in other words, trees [26] and their variants. This topological preference is definitely due to the fact that no two adjacent nodes can carry alike charge. A few examples of four-membered closed rings either isolated or ‘fused along an edge’ [26] were also found. For closed rings, the topological constraints due to charge allows only an even number of nodes.

**Figure 3.**
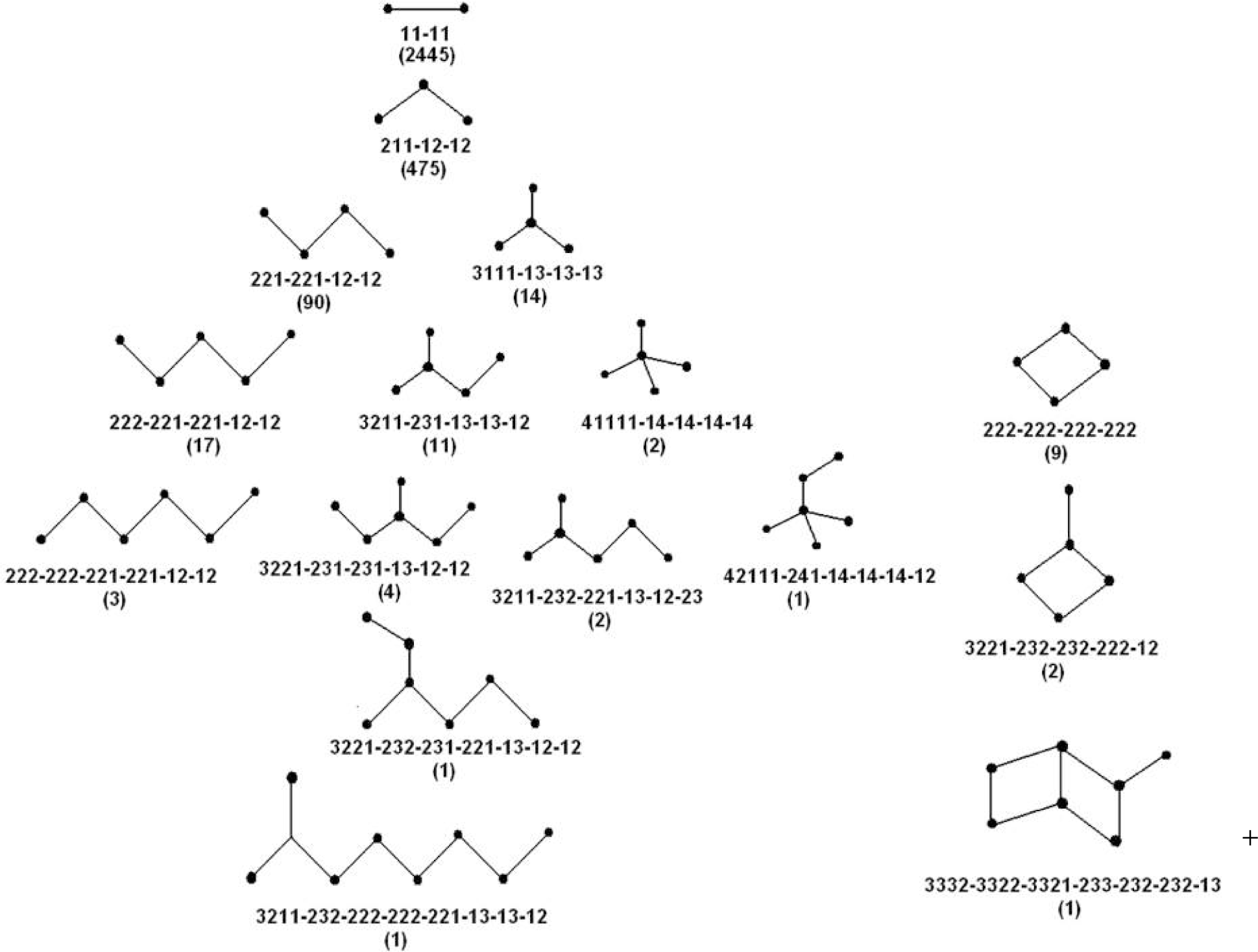
Statistical distribution of networks of ionic bonds within proteins. Each network topology is demonstrated by the motif identifier (numeric string). The number of such networks found is given in parenthesis below the identifier.

### Electrostatic Complementarity

Overall, a mild enhancement in *E_m_* was observed for charged residues, involved in salt-bridges **(Fig.4)**, with the exception of histidine which was found to prefer metal coordination sites more than salt-bridges [26]. The highest average value of 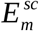 was obtained for Glutamate (0.68) which also had the highest increment in <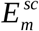> upon inclusion into a salt-bridge. Arginine exhibited the highest propensity to form ionic bonds (5.83), compared to other charged residues (Glu: 4.77, Asp: 3.92, Lys: 3.43). The participation of histidine in ionic bond networks was by and large negligible (propensity: 0.22).

**Figure 4.**
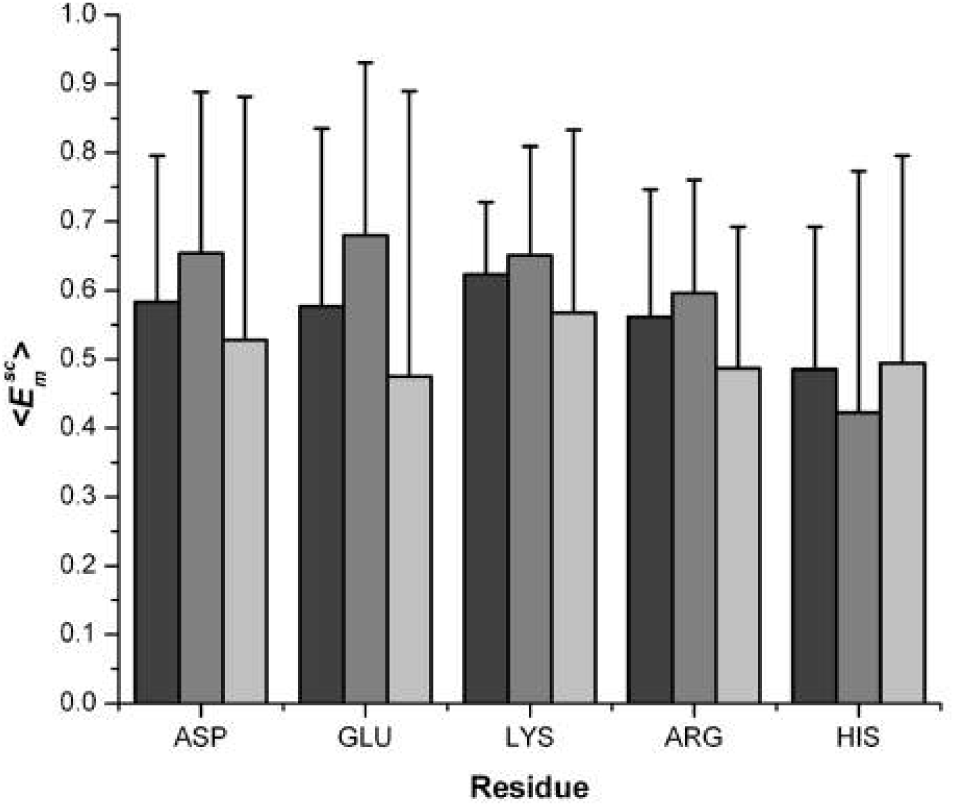
Charged residues involved in salt-bridges lead to only mild enhancement in. The figure shows mean values of (filled thick bars) along with their standard deviations (thin error bars) for charged residues involved in salt-bridges (gray), not involved in salt-bridges (light gray) and pooled together (deep gray). As can be seen, histidine (positively charged) shows a reverse trend.

### Compositional and Geometrical preferences

Several instances have been recorded where bifurcated salt-bridges contribute more to the electrostatic stabilization within proteins than the isolated ionic bonds [10], [13], [14]. With this emphasis, bifurcated salt-bridges (**Fig.5**) were further analyzed for compositional and geometrical bias (**Table.1**). Compositional preferences were clearly distinguishable for arginine containing salt-bridges (75.4% of the whole set) with Glu-Arg-Glu having the highest occupancy (17%). Similar preferences have been previously observed for arginine-glutamate salt-bridges in case of helix stability [10].

**Figure 5.**
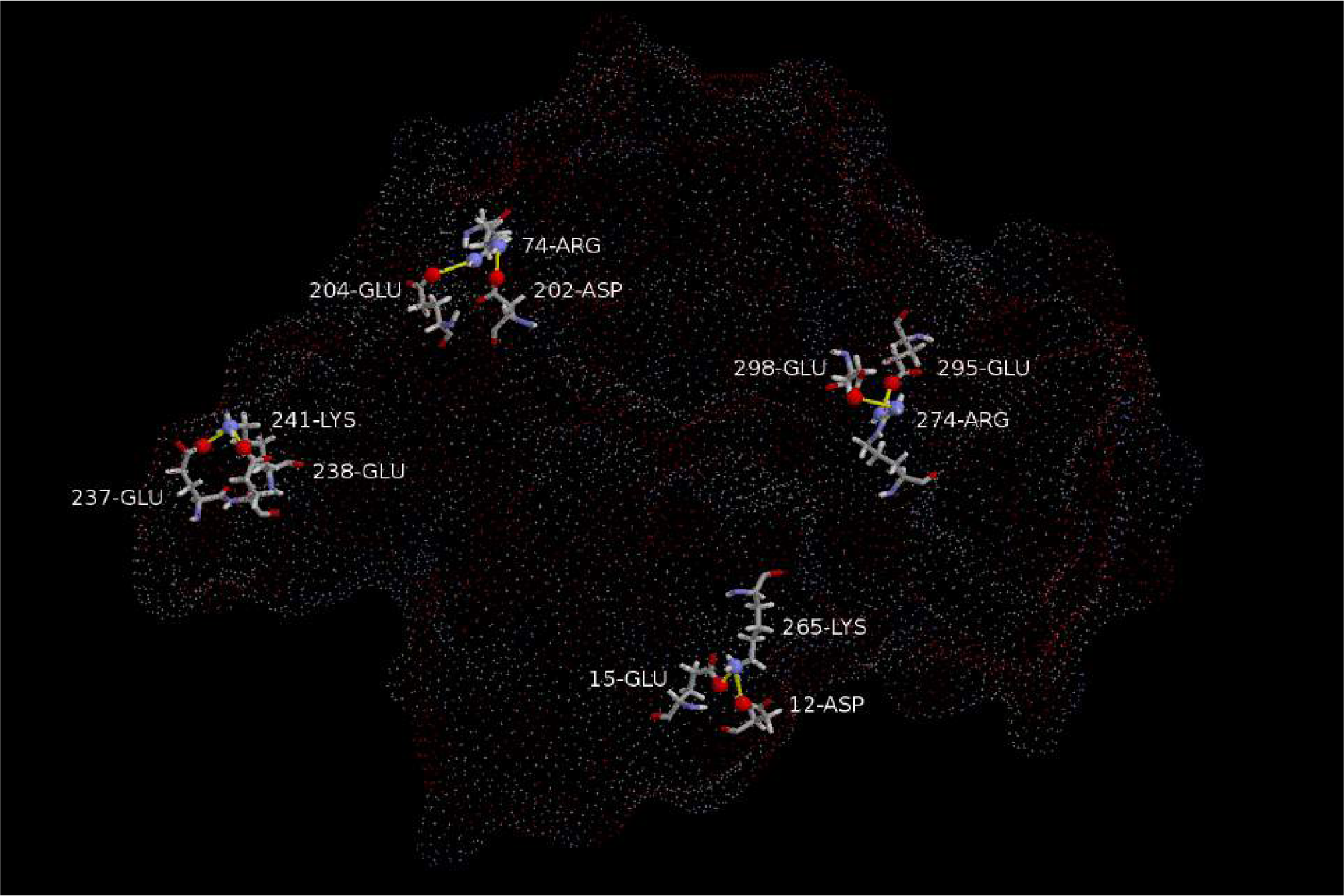
Bifurcated salt-bridges within proteins. All bifurcated salt-briges identified by the present algorithm in the protein 3B8X. The salt-bridges have been displayed as yellow sticks joining red balls (charged atoms) shown over the molecular dot surface of the protein. The molecular surface was constructed by EDTsurf [42]. Most amino acids involved in the salt-bridges are partially exposed to the solvent. The examples include both catatonic (three of them) as well as anionic centers (one) and instances of both short (three of them) and long-range contacts (one). Figure created using Rasmol (http://www.openrasmol.org/).

**Table 1.**
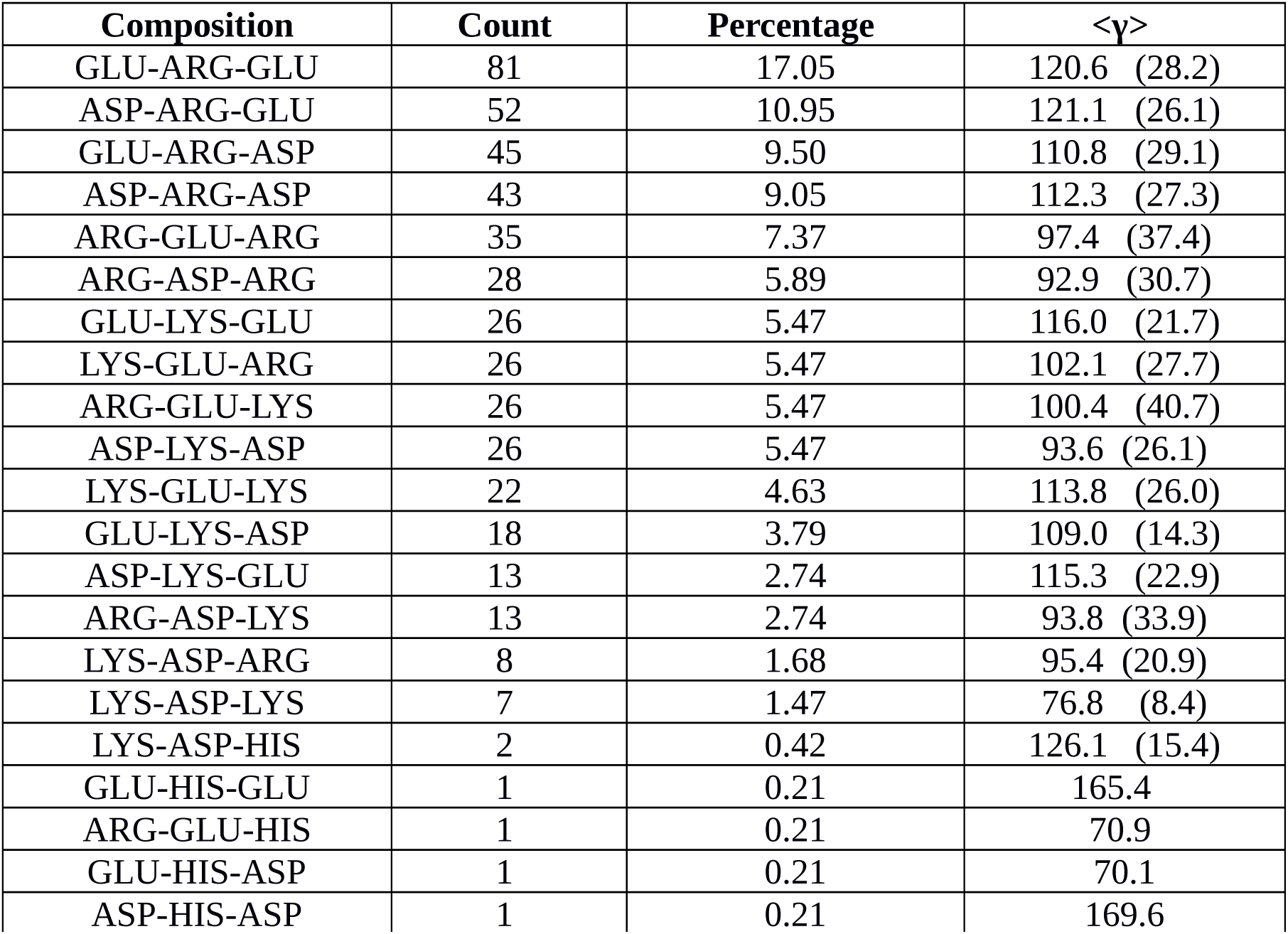
Composition and Geometry of the Bifurcation angle (Y)

| Composition | Count | Percentage | $\langle\gamma\rangle$ |
| --- | --- | --- | --- |
| GLU-ARG-GLU | 81 | 17.05 | 120.6 (28.2) |
| ASP-ARG-GLU | 52 | 10.95 | 121.1 (26.1) |
| GLU-ARG-ASP | 45 | 9.50 | 110.8 (29.1) |
| ASP-ARG-ASP | 43 | 9.05 | 112.3 (27.3) |
| ARG-GLU-ARG | 35 | 7.37 | 97.4 (37.4) |
| ARG-ASP-ARG | 28 | 5.89 | 92.9 (30.7) |
| GLU-LYS-GLU | 26 | 5.47 | 116.0 (21.7) |
| LYS-GLU-ARG | 26 | 5.47 | 102.1 (27.7) |
| ARG-GLU-LYS | 26 | 5.47 | 100.4 (40.7) |
| ASP-LYS-ASP | 26 | 5.47 | 93.6 (26.1) |
| LYS-GLU-LYS | 22 | 4.63 | 113.8 (26.0) |
| GLU-LYS-ASP | 18 | 3.79 | 109.0 (14.3) |
| ASP-LYS-GLU | 13 | 2.74 | 115.3 (22.9) |
| ARG-ASP-LYS | 13 | 2.74 | 93.8 (33.9) |
| LYS-ASP-ARG | 8 | 1.68 | 95.4 (20.9) |
| LYS-ASP-LYS | 7 | 1.47 | 76.8 (8.4) |
| LYS-ASP-HIS | 2 | 0.42 | 126.1 (15.4) |
| GLU-HIS-GLU | 1 | 0.21 | 165.4 |
| ARG-GLU-HIS | 1 | 0.21 | 70.9 |
| GLU-HIS-ASP | 1 | 0.21 | 70.1 |
| ASP-HIS-ASP | 1 | 0.21 | 169.6 |

The angle subtended by three residues forming a bifurcated salt-bridge was computed as follows: except for lysine, which has an unique charged nitrogen atom (NZ), the effective or resultant charge centers were determined as the midpoint of the two degenerate charged O (aspartate, glutamate) and N (arginine, positively charged histidine) atoms. The bifurcation angle (γ) between the two vectors connecting the three charge-centers was then computed, which was found to be obtuse and fairly well constrained (∼110° ± 30°) irrespective of the residue composition.

### Contact Order

Contact order (co) of the interacting pairs of ions involved in a salt-bridge was calculated as the separation of the two amino acid residues in sequence space divided by the full length of the polypeptide chain. The majority of the interactions were found to be short-range contacts with (∼55% hitting a value lesser than 0.1 which physically means that the ioinic bonds formed between amino acid residues which are within 10% of each other in sequence space with respect to the whole length of the polypeptide chain. The overall distribution follows an asymptotic decay for long-range contacts characterized by a long tail and a median of 0.079 (**Fig.6**). Again, significant preferences were further observed for a co value of 0.03 accounting for 42% of the whole population of salt-bridges - which is consistent with the earlier observations [8] that salt-bridges are not only preferably formed between residues close than distant in sequence space but also at specific small sequence separations. No major preference was observed for ionic bonds involved in bifurcated salt-bridges compared to salt-bridges in general.

**Figure 6.**
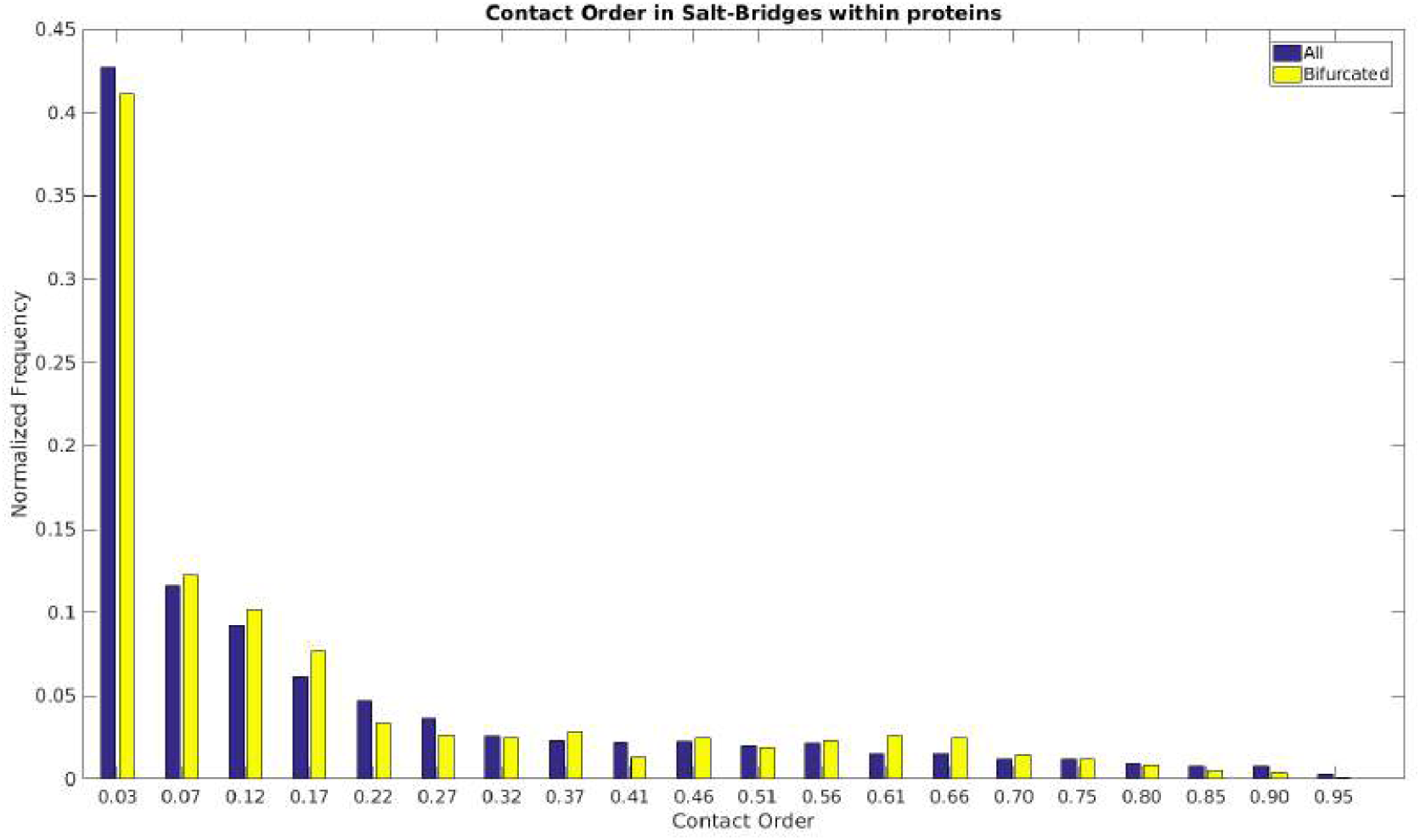
Contact Order in Salt-bridges within proteins. The bar diagram plots contact order (see **Materials and Methods**) for inter-residue contacts found in salt-bridge networks in general. As evident from the plot, most contacts are short-ranged and no particular preference could be found in bifurcated salt-bridges compared to salt-bridges in general.

### Bifurcated salt-bridges in protein-protein / protein-ligand interactions

There exist a very fine line of difference between folding and binding of proteins, and the gap is even narrower if viewed from the common conceptual platform based on complementarity. More specifically, it was demonstrated in a previous study [6] that folding could actually be envisaged as the docking of interior components onto the native polypeptide chain upon folding. From that view, there is practically very little or no difference between salt-bridges involved in the structural stability of a folded protein and those involved in protein-protein interactions (binding) in terms their physico-chemical attributes. However, as a subject of an individual study, it was of interest to know the role of bifurcated salt-bridges in triggering and stabilizing inter-protein associations. In other words, the study could not have been complete without a thorough statistical analysis involving bifurcated salt-bridges in (i) protein-ligand and (ii) protein-protein interactions. The first calculation was performed in the same ‘globular protein’ database while another database of native protein-protein complexes containing 1879 high resolution crystal structures were assembled for the second (see **Materials and Methods**).

For the first calculation, only those proteins from the globular protein database were gathered which contained any non-protein (hetero) atoms other than the crystallographic water molecules or buffer components. The subset of structures involving one or more bifurcated salt-bridges which were in contact with any of the hetero atoms were further identified. An atomic separation of maximum 4 Å was allowed between any heavy atom coming from the bifurcated salt-bridge and any of the nonprotein atoms to filter out the contacts. This stringent contact criteria was set to keep uniformity with the adapted definition of salt-bridges and also to remove crystallographic artifacts of pseudo-contacts. A total of 69 structures were obtained where one or more bifurcated salt-bridge was directly involved in ligand binding, adding upto a total of 92 salt-bridge-ligand interactions (**Supplementary Table. S1**). The ligands range from isolated metal ions to bulky prosthetic groups, and from essential co-factors to coenzymes and enzyme-substrates (**Fig.7**).

**Figure 7.**
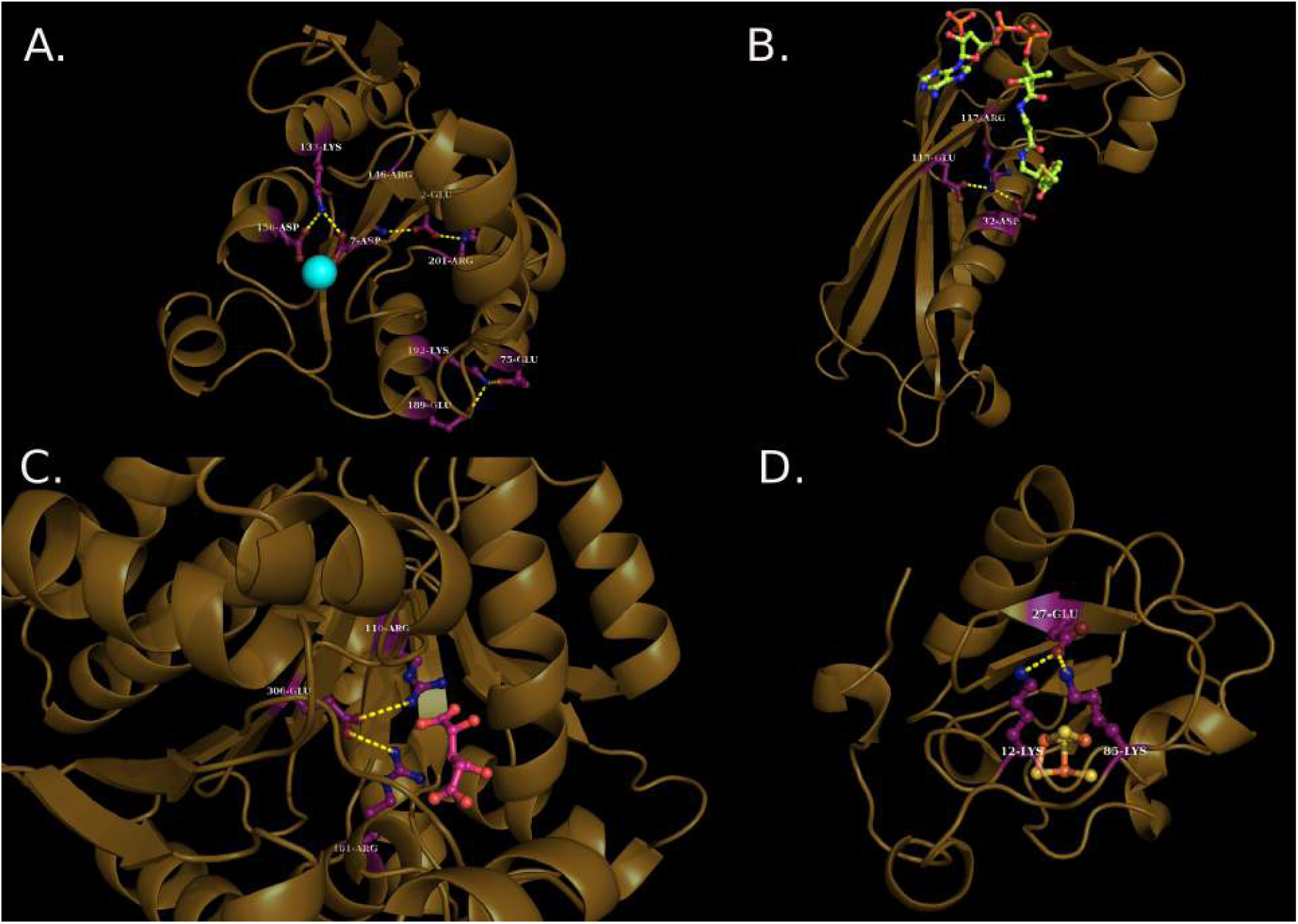
Bifurcated salt-bridges anchoring ligands within proteins. A. Homoserine Kinase from *pseudomonas aeruginosa* (1RKU) have three bifurcated salt-bridges, all of which are involved in the packing of secondary structural elements (Helices and Sheets). Moreover, one of them (7-ASP∼133-LYS∼156-ASP) is additionally coordinating a Mg+2 ion, mediated by the e-NH3+ group middle Lysine. **B.** Isocitrate dehydrogenase (1LWD) from porcine heart mitochondria have a centrally located bifurcated salt-bridge (101-ARG∼306-GLU∼110-ARG) which binds the reaction-substrate, Isocitrate by two ionic bonds mediated by the guanidium groups of two Arginines with two carboxylic acid groups coming from the isocitrate. This is an unique example of formation of a closed quadrangle protein-substrate salt-bridge initiated by a bifurcated salt-bridge. **C.** Hydrolase from *pseudomonas sp.* (1LO7) have a central bifurcated salt-bridge (32-ASP∼117-ARG∼115-GLU) which anchors to the substrate-coenzyme / extended prosthetic group, 4-Hydroxybenzoyl CoA via an ionic bond mediated by an Aspertate side-chain. **D.** Ferridoxin (1PC4) from *azotobacter vinelandii* have a peripheral bifurcated salt-bridge (85-LYS∼27-GLU∼12-LYS) which helps the protein to its cofactor, the iron-supher cluster (Fe3S3).

A classic and interesting example was found in the binding of Isocitrate Dehydrogenase (PDB ID: 1LWD) to its substrate, isocitrate, wherein, guanidium groups from two arginines were involved in the formation of two further ionic bonds with the carboxylic acid groups coming from the isocitrate (**Fig.7B**). In effect, the topology of the full interactome becomes a closed quadrangle (222-222-222-222) from an open triple / angle (211-12-12: a bifurcated salt-bridge) considering the isocitrate also to be part of the extended protein-substrate salt-bridge network. Point to note is that the bifurcated open triple only requires one of its member (essentially the middle node in the network) to involve in two simultaneous interactions whereas the quadruplet requires every member to have two extended hands in order to make it closed. The example also stands out to be interesting, since, not only the initial salt-bridge is bifurcated (or fork-like), but also, all the key interacting chemical groups are bifurcated themselves.

The second calculation involving protein-protein interactions is perhaps a more direct and clear evidence of the role played by bifurcated salt-bridges in protein functions. Indeed, some of the compositional and geometric features of protein-protein interfaces are quite odd and in significant contrast to that of the interior. For example, isolated nonpolar residues are often found at protein-protein interfaces, enclosed by polar or charged amino acids [31], in contrast to the hydrophobic clusters found regularly within protein interiors. These contrasting features make it more interesting to investigate the distribution and functional specificity of salt-bridges at the interface relative to the interior. Only those salt-bridges were considered, which involved both the molecular partners (i.e., the two interacting polypeptide chains). In other words, in all the interfacial salt-bridges extracted, both the interacting polypeptide chains contributed at least one salt-bridge forming residue. Again, but for the trivial case of isolated salt-bridges, bifurcated salt-bridges were found in the largest majority (among all other networks) and prevalent at the interface. Out of a total of 1879 native PPI complexes, 1088 contained one or more salt-bridges directly at the interface. Again, 211 complexes were found to contain one or more bifurcated salt-bridges (**Supplementary Table S2**) at the interface adding upto a total of 283 bifurcated salt-bridges (211-12-12), the second highest after the isolated salt-bridges (11-11: 2022).

In several instances, there were actually multiple salt-bridges at the protein-protein interface, with a variety of relative spacial separation among them. In some cases, for example, the interfacial salt-bridges occurred in such close proximity (**Fig.8A**) that is reminiscent of a potential cooperative effect in order to promote ioinic interactions at the interface, while, in other cases, the multiple salt-bridges were distantly spaced spanning a large contour of the interface (**Fig.8B**) as if to cover more surface area buried upon complexation. In character, the interfacial bifurcated salt-bridges resembled molecular clips between the interacting protein surfaces.

**Figure 8.**
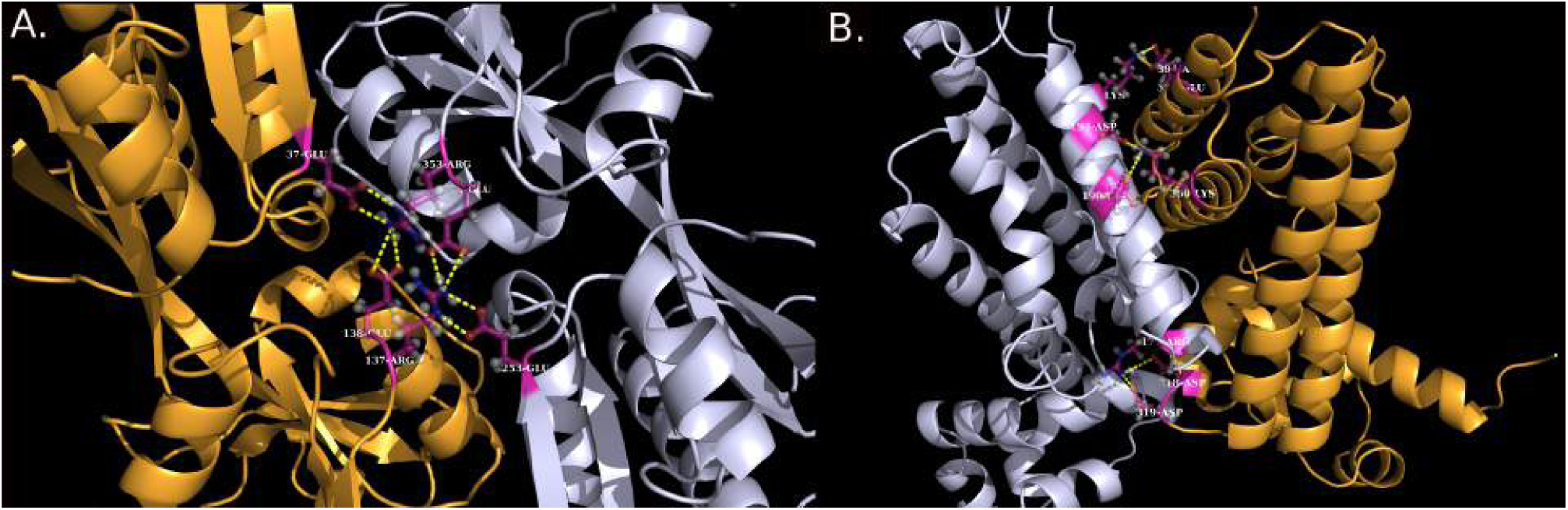
Bifurcated salt-bridges clipping protein-protein interfaces. A. Multiple bifurcated salt-bridges (253-GLU-B∼137-ARG-A∼354-GLU-B; 138-GLU-A∼;353-ARG-B∼;37-GLU-A) centrally located at the interface acting in cooperation to clip the interface in an effector binding domain of a Benm Variant (2H9B). **B.** Multiple bifurcated salt-bridges (390-GLU-B~160-LYS-A∼;394-ASP-B; 319-ASP-B∼;177-ARG-A∼;318-ASP-B; 194-ASP-A∼;360-LYS-B∼;190-GLU-A) clipping the interface from distant locations while covering a large surface area in a transcriptional regulator, TetR family protein complex from *Cytophaga hutchinsonii* (3EUP).

### Salt-bridge mediated interactions between ordered and disordered regions in partially disordered proteins

Intrinsically disordered proteins have served an intense subject in protein-science especially in the last decade. A significant fraction of the effort has been devoted towards deciphering a possible disorder code and unraveling the physical chemistry associated with the disorder-to-order transition that the residues (namely, ‘protean’) undergo upon binding [18], [32], [33]. Machine learning have been used vigorously in the sequence-based predictions of both the disordered as well as the protean residues [34], [35]. It has been an established fact by now that the disordered regions are rich in charged and polar amino acids in contrast to having a low hydrophobic content [36] – serving as plausible mechanistic insights into the origin of their disorder. It has also been suggested that the associated low hydrophobic content disfavors self-folding [37] by potentially decreasing the number of possible two-body contacts [38].

Unlike the nucleation-condensation model reminiscent of the hydrophobic packing [26] and folding [39], [40] for globular proteins, the aforementioned features practically rule out the possibility of hydrophobic interactions playing an equivalent determining role in the partial folding of the IDPRs, rather, emphasizes on the contribution of ionic bonds and atomic contacts mediated by charged and polar residues being more influential. To test this hypothesis, proteins containing structural disorder at an amount greater than 50% [18] were assembled and the missing disordered regions modeled using MODELLER (see **Materials and Methods**). The relative percentage fraction of secondary structural content (calculated by Stride) had an overwhelming preference for coils (or disordered loops) in these modeled structures, as expected. The fraction of coils ranged from 12.3% to 96% with an average of 61.75% (± 18.17%) which was accompanied by an average fraction of 16.02% (± 10.33%) turns.

It is obvious that the modeled structures only represent a reduced conformational space out of the whole plethora of possible conformational ensembles attributed to the disordered regions. However, addressing the issue of selecting the best structural representative should involve time-average structures subjected to large-scale molecular dynamic simulations falling outside the scope of the current study. Yet, it was important to identify the extreme cases of largely asymmetric protein models with completely unstructured long wobbling loops present even in the selected best model (out of the 10 top models) and eliminate them from the current atomic-contact-based calculation to remove statistical artifacts. The long wobbling disordered loops in these models occupied a huge volume of physical dimension with practically no chance of participating in internal packing - which was confirmed from their individual visual inspection in Pymol (**Supplementary Fig.S1**). Especially since the objective was to study the role of salt-bridge mediated atomic contacts in the partial folding of these proteins, it was necessary to remove these outliers. The pseudo-centroid of the protein molecule was calculated from the collection of experimentally solved atoms alone and the distance of each completely exposed atom (see **Materials and Methods**) was calculated from this pseudo-centroid. The root mean square deviation of these distances was chosen as the parameter (igA) indicative of globular asymmetry and the largely asymmetric models were discarded based on an *ad-hoc* cutoff set to this parameter. The same parameter, when calculated in the database of globular proteins (see **Materials and Methods**) gave a modest value of 4.92 (± 2.2) reflecting their approximately spherical shapes, but for a few elongated proteins. In drastic contrast, the average igA value obtained for the disordered proteins was 29.35 (±30.34) with a maximum of 171.37 reflecting significant asymmetry and no sign of even partial folding given the volume constraints reminiscent of a crowded living cell. We surely did not want the calculation to be contaminated by such unreal atomic models. Given all these, it still remains largely empirical, non-autonomous and context dependent to set an appropriate cutoff given the vastly unknown system of IDPs present significant contrast to that of the globular protein system. Thus, visual intervention appeared indispensable coupled with an arguable statistical rationale. One way could be to retain only those models for which the average igA value was no more than μ +3σ with respect to the values obtained from the globular proteins (μ: mean, σ: standard deviation) concomitantly allowing about a two-fold increase in the associated standard deviation. A cutoff (in igA) of 20 Å gave rise to an average igA of 11.5 (± 4.1) which was found to be the closest match to the above filtering criteria, reducing the number of structures to 62 (a ∼57% recovery). To note is the fact that even this cutoff should be considered fairly relaxed considering the objective of not to miss out on any potential candidate giving rise to enough disorder-order atomic contacts. This very method (using a range of cutoffs for igA) was also used previously [6] to successfully standardize and delineate the boundaries between different grid size bins and thereby effectively fix the box dimension (cubic lattice) for globular proteins during the course of their continuum electrostatic calculations by the Poisson Boltzmann method using Delphi [30]. The correctness of the filtering criteria was also cross-validated by the use of the Accessibility score (see **Materials and Methods**) which was raised from 0.0018 (± 0.042) for the unfiltered set to 0.0033 (± 0.048) for the filtered set, rendering roughly a ∼2-fold increase in the average score with standard deviations falling in the same range. Therefore, the filtered set of disordered protein models does appear to be better in terms of solution stability and hence more realistic.

To test whether salt-bridge mediated interactions were influential in the partial folding of these IDPRs, salt-bridges were identified following the same consistent criteria (see **Materials and Methods**) from the whole molecules for each protein. Other atomic contacts were then identified between any pair of heavy atoms within 4 Å of each-other where one heavy atom was contributed by a residue located in the modeled disordered regions and the other from a charged residues involved in a salt-bridge irrespective of its location on the protein. A total of 180 salt-bridges were identified contributing to 4630 salt-bridge mediated non-salt-bridge atomic contacts leading to 383 inter-residue interactions. Again, isolated ionic bonds had the largest frequency (11-11: 108) followed by bifurcated salt-bridges (211-12-12: 12). The salt-bridge mediated contacts, by definition, should involve both polar (or charged) and non-polar atoms coming from either the main-chain or the side-chain of the interacting residues. The distribution of contacts showed a large preference towards main-chain atoms constituting ∼80% of the atoms involved in contact (main-chain ∼ main-chain contacts: 2962, main-chain ∼ side-chain: 1441 and side-chain ∼ side-chain: 227) which is most likely due to the fact that the (few) charged side-chain atoms (-NH+ and −COO-groups) are already occupied in salt-bridges and therefore somewhat less available to forming other contacts compared to the main-chain atoms. The contacts mediated by slat-bridge forming charged residues indeed appeared to be subservient in the partial folding of these quasi-stable proteins. Several prototype interaction modes were identified (**Fig.9**) spanning from bridging loops (**Fig.9.A**) bending and turning around of loops (**Fig.9.B-C, G**), helix-loop clippings (**Fig.9.C-H**), partial stabilization of loops by promoting a series of contacts (**Fig.9.C, F, G**), bringing helices together by mediating contacts in helix-loop-helix and other related secondary structural motifs (**Fig.9.I-J**) etc. The results should guide to explore the modus operandi of these partially disordered proteins - subject to future studies.

**Figure 9.**
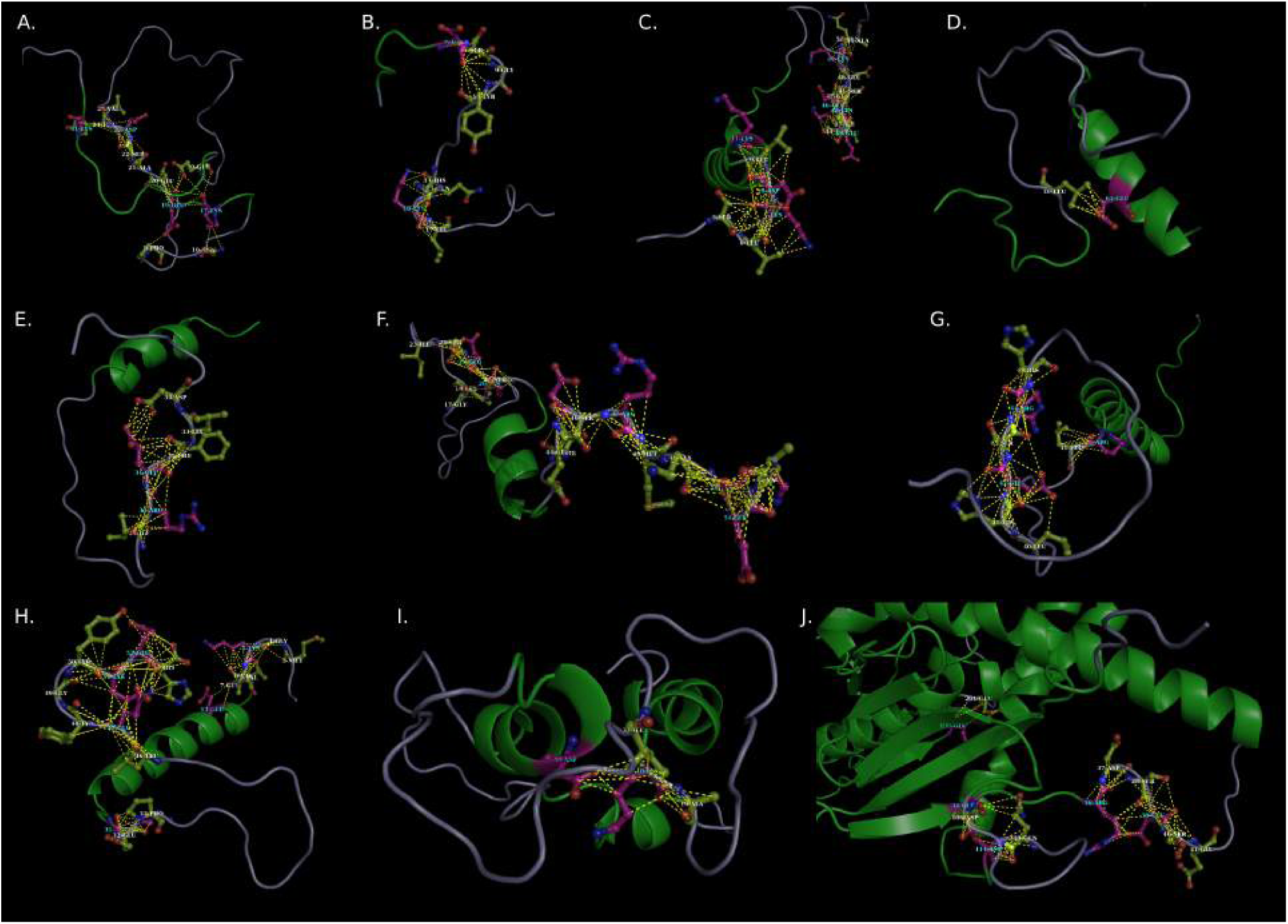
Salt-bridge mediated contacts in partially disordered proteins. The figure presents selected examples of salt-bridge mediated interactions representing a variety of interaction modes between residues located in disordered loops and salt-bridge residues located in the structured regions in modeled structures containing IDPRs. A vivid description of the different modes of interaction have been elaborated in the corresponding section in the main text. The PDB ID’s are as follows. (A) 3KND (B) 3IFN (C) 3SJH (D) 3MN7 (E) 2JZ3 (F) 4A1G (G) 4G91 (H) 1H8B (I) 1FH1 (J) 4H62. Residue numbers in each modeled structure commence from 1 following the SEQRES records (see main text) in the corresponding PDB file and all salt-bridges are labeled in that way.

### Salt-bridge mediated interactions in stabilizing disordered protein regions upon binding

We also had a close look on a few most popular IDPs / IDPRs for which there are available experimental structure either for in their free form or in their bound form. The motif was to investigate whether their structural stabilization in the bound form or the lack of stabilization in the free form had any influence of salt-bridge mediated interactions. Due to their intrinsic disorder, IDP’s generally does not form stable and diffractable crystals in their free forms. And even, in their bound forms, only fragments of the whole protein are all that are available. These fragments representing the ‘disorder-to-order’ transitioning residues upon binding are categorically termed ‘protean’ (mutable) fragments and a growing field of computational tools are being developed for their sequence based predictions [34], [35].

### Case Studies on Tau

One of the most popular IDP is surely the Tau protein which is one of the prime causal factors for neurofibrillary degeneration in the Alzimer’s disease. Apart from having a solution (NMR) structure in its free form (2N4R), there are a few crystal structures available for this protein, bound to different cellular partners and also to diagnostic markers, most of which however only covers a small fraction (3-15 residues) of the whole protein which is 202 amino acids long. Among the available complexes, the current analyses opted for 4TQE, 5DMG and 5BTV, where, in the first two, the Tau-peptide was bound to anti-Tau antibody and in the third case, a different peptide fragment of the same protein is bound to epithelial cell marker protein 1, stratifin forming the sigma complex. The study also included the NMR structure (the first and the lowest energy model) of the standalone molecule into the analyses. Detail structural investigation from the perspective of salt-bridge mediated stabilization did bring out some really interesting features.

Tau, in its free form (i.e., the 202 residue long fragment) is partially disordered containing roughly 50% of beta-sheet and ∼50% of coil, as revealed from the NMR structure. The structure was found to have 4 isolated salt-bridges (11-11) all of them playing pivotal roles in its folding, by either getting involved in loop-closure or joining distant secondary structural elements in space (**Fig.10.A**). Two of the salt-bridges were located on extreme opposites along a horizontal pseudo-axis, stabilizing (intra-and inter-strand) turns whereas the other two salt-bridges were closely spaced, aligned orthogonality to one-another, forming a T-shaped pair. The relative orientation of the stem of the ‘T’ is vertical with respect to the aforementioned horizontal pseudo-axis, giving mechanical support to the overall structure. In effect the roof of the ‘T’ resembles a fly-over.

**Figure 10.**
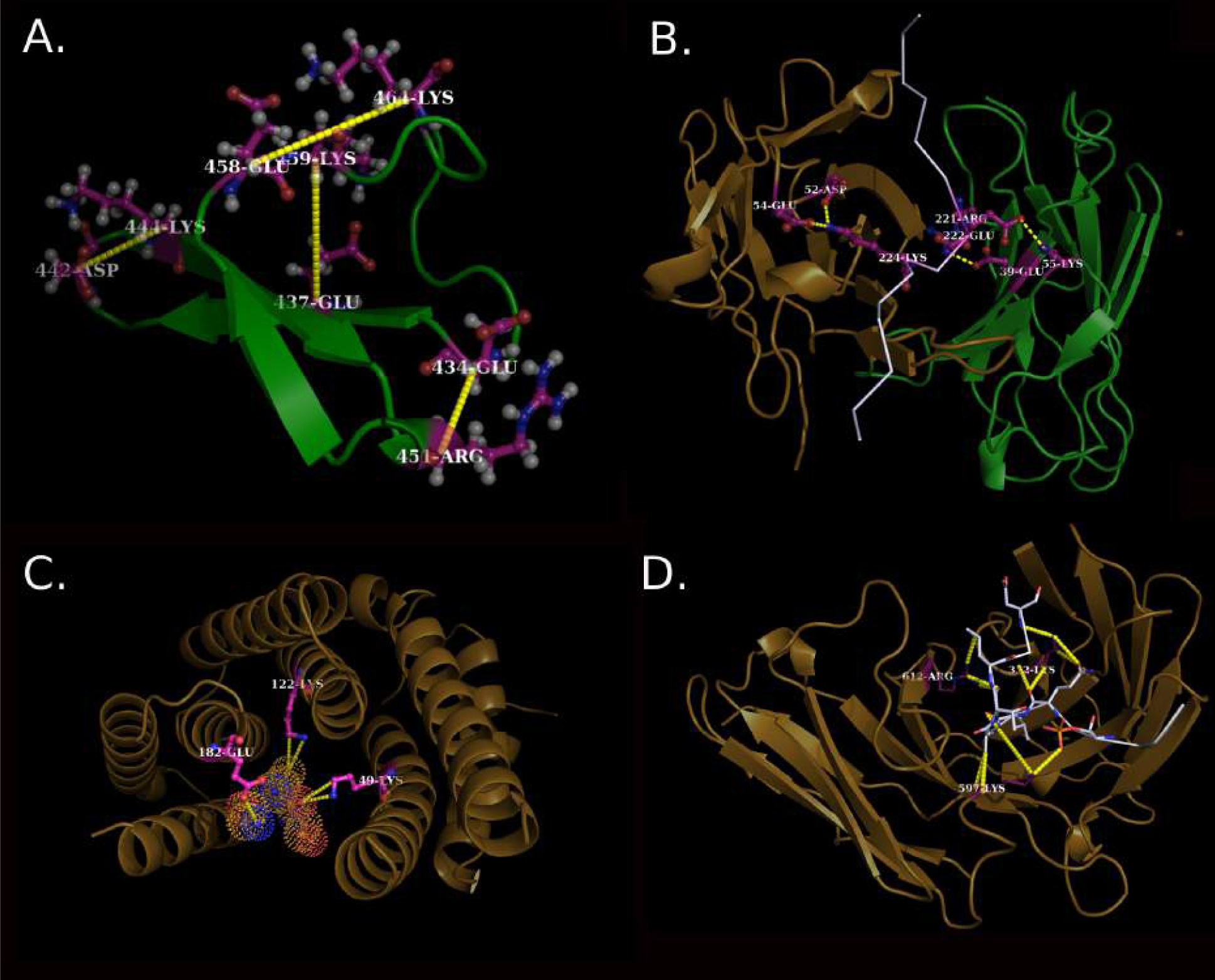
Role of salt-bridges and salt-bridge mediated interactions in the formation of the Tau fold and in the stability of different Tau peptides upon binding. Panel (A) presents the NMR structure of the 202 amino acid long fragment of the Tau protein in its free form (PDB ID: 2N4R) whereas (B) (C) and (D) presents different Tau peptides in their bound forms (PDB IDs: (B) 4TQE, (C) 5DMG (D) 5BTV). (A) The two orthogonaly oriented salt-bridges form together a T-shaped sub-structure in the free form of Tau which appear to be instrumental in holding the fold. (B) One bifurcated and two isolated salt-bridges stitches the Tau peptide with the anti-Tau antibody. (C) and (D) presents cases where the Tau peptides (different to that of the one in A) are stabilized upon binding to their partners not only directly involving salt-bridges but also promoting salt-bridge mediated interactions. A more detail elaboration is given in the main text.

In first of the complexes, namely, 4TQE, a lysine (K224-Tau) situated centrally in the thread-like 15 amino acid long Tau peptide was stitched meticulously by two negatively charged residues (D52-H, E54-H) coming from the heavy chain (H) of the anti-Tau antibody, forming an interfacial bifurcated salt-bridge (**Fig.10.B**). In addition, there are two more isolated salt-bridges (K55-L ∼ E222-Tau, E39-L ∼ R221-Tau) found at the interface, contributed by the light chain (L), stitching the disordered peptide thread even further along the deep groove of the antibody Fab fragment. Important to note that the charged residues involved in the salt-bridges are closely spaced and located centrally on the peptide and thereby enhancing the strength and stability of the binding. The role of ionic bond stabilization in this ‘disorder-to-order’ transition thus can not be undermined. In both of the other two examples (5BTV, 5DMG), charged residues seemed to be crucial in the binding and stabilization of the peptides. Important to note is that all three Tau peptides studied were different and non-overlapping. In more detail, ionic bonds as well as N…C non-covalent contacts coming from charged residues from both the Tau peptide and its molecular partner were found to be instrumental in its stabilization (**Fig.10.C-D**). The inter-molecular ionic bonds, in effect, forms extended slat-bridge networks in conjugation with one-another.

### Case study on Crambin

Another case study was executed on the plant protein Crambin (from *Crambe hispanica*) solved at an ultra-high resolution (0.54 Å) (PDB ID: 1EJG) presenting an unique case of a protein having multiple occupancy atoms throughout the whole chain with each isoform being individually disordered (http://deposit.rcsb.org/format-faq-v1.html). This protein was later solved at an even higher resolution (PDB ID: 3NIR, 0.48 Å) and both the ultra-high resolution structures served as great candidates in the study of protein helices reconciled with three-centered hydrogen bonds [41]. Interestingly, the whole protein (1EJG) gave rise to just a single isolated salt-bridge (17-ARG∼23-GLU) bringing together the largest helix and a long irregular loop. Most remarkably the two charged residues (17-ARG, 23-GLU) involved in this single isolated salt-bridge were further involved in 113 more atomic contacts involving 21 other residues in a manner to hold the entire protein fold like that of a bridge (**Fig.11.A**). The atomic contacts were evenly distributed among non-polar (C: 56) and polar atoms (O, N, S: 57). The act of retaining the fold together was especially eminent towards the more structured half of the fold containing the two helices. Therefore, the so-called ‘isolated’ salt-bridge does not appear to be isolated in a broader structural sense, rather instrumental in the integrity of the entire fold. The results were cross-validated in the higher-resolution structure, 3NIR (**Fig.11.B**).

**Figure 11.**
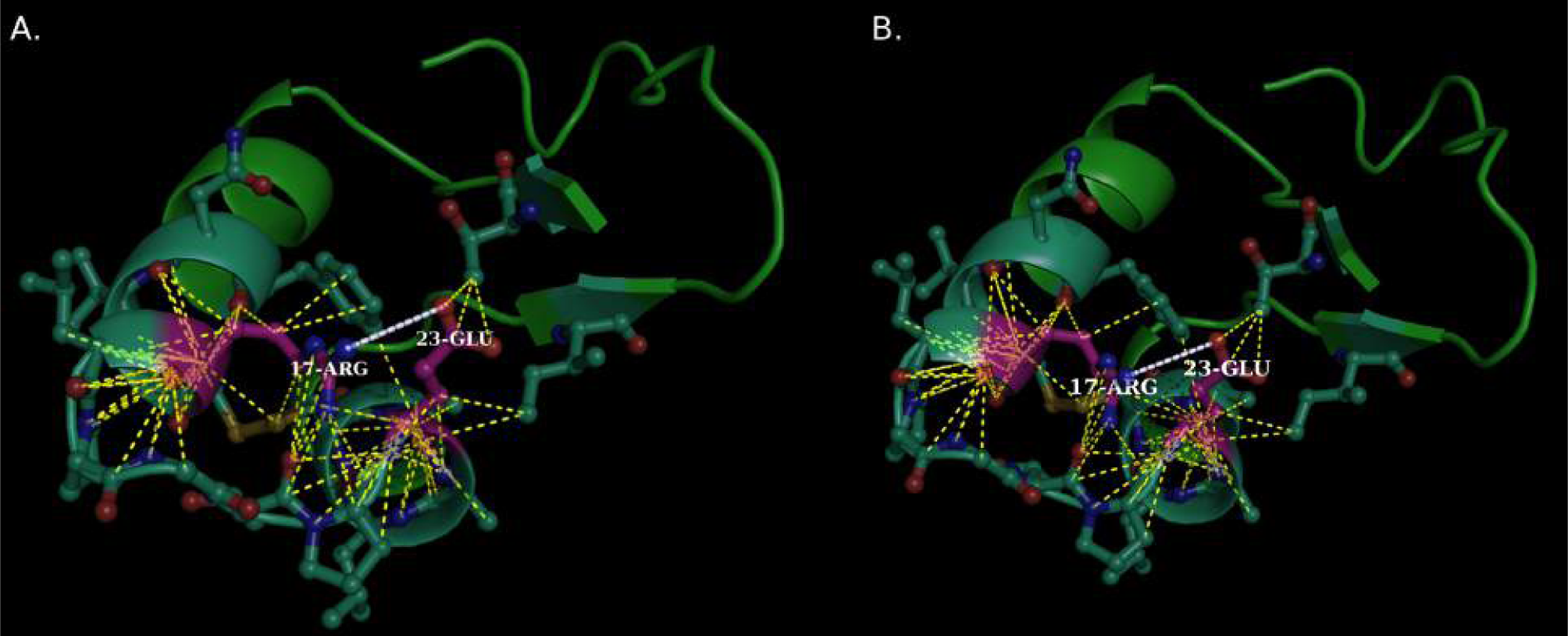
Structural integration and stabilization of Crambin involving salt-bridge mediated interactions. (A) and (B) presents two ultra-high resolution structures of the protein Crambin (PDB ID: 1EJG, 3NIR respectively) solved at gradually increasing resolutions and show identical results. Both structures were used in the study as a mean to cross-validate each other. As elaborated in detail in the corresponding main text, the structures represent an unique case of a single salt-bridge (17- ARG ∼ 23-GLU) promoting more than 100 atomic contacts involving more than 20 other residues which is instrumental in the structural integration of the protein fold.

## Conclusion

The current study analyzes the variety and extent of different modes of association (networks) pertaining to salt-bridges on or within folded proteins and at protein-protein interfaces. The ‘bifurcated salt-bridge’ appear to be a special and prevalent salt-bridge motif. Apart from serving a major component in the meticulous electrostatic balance attributed to a folded protein, a detailed functional characterization of salt-bridges show their key participation in binding ligands, cofactors, metal ions, prosthetic groups and also in promoting a variety of protein-protein interactions. One very interesting functional motif involving bifurcated interfacial salt-bridges could be envisaged as molecular clips stitching large surface contours of interacting protein-protein interfaces. Salt-bridge mediated interactions seem pivotal in the partial folding of proteins containing large amount of disordered regions. The results should benefit in the conceptualization of how these proteins manage to keep necessary amount of disorder even in their functionally active bound forms. Lastly, the current study should be beneficial towards the proposed notion of geometrically specific designable interactions involving salt-bridges. Though, it is much difficult to control the conformational space of the side-chain rotamers for elongated charged residues (Lys, Arg, Glu) [8], molecular dynamics simulation studies have elucidated the role of configurational entropy of salt-bridge networks in thermostability of peptides [9], and, the current study should provide important rules of thumb, instrumental in further research down that line.

## Acknowledgment and Funding

We express our heartfelt gratitude to Prof. Anjan Kumar Dasgupta (Department of Biochemsitry, University of Calcutta) for his constant support and encouragement for the work. The work was supported by the Department of Science and Technology – Science and Engineering Research Board (DST-SERB research grant **PDF/2015/001079)**.

